# Neural Timescale of Adolescents Major Depressive Disorder

**DOI:** 10.1101/2025.07.01.662431

**Authors:** Yujun Gao, Sanwang Wang, Hanliang Wei, Jinzhi He, Geng Liu, Lang Liu, Xiaoqiang Liu, Xiao Li, Xianwei Guo, Xiaobo Liu

## Abstract

Adolescent major depressive disorder (MDD) is characterized by heterogeneous symptomatology and complex neurodevelopmental underpinnings. Here, we investigated whether cortical intrinsic neural timescales (INT)—a measure of temporal stability in neural activity—are altered in adolescents with MDD and whether these alterations relate to clinical symptoms, suicidality, early-life adversity, and underlying neurobiological mechanisms. Using resting-state fMRI in adolescents with MDD and healthy controls (HCs), we found widespread reductions in INT in frontal, parietal, and sensorimotor regions, with a notable prolongation in the left temporoparietal junction. Disrupted timescales significantly impaired network modularity and clustering, and SVR-based machine learning revealed that altered INT patterns predicted individual depression and anxiety severity. INT abnormalities were further associated with suicidal ideation and childhood trauma, particularly emotional and physical neglect. Biophysical modeling linked INT variations to local recurrent excitation and external input strength, differing between HCs and MDD. Spatial correlations with PET-derived neurotransmitter receptor maps demonstrated that INT alterations colocalize with serotonergic, dopaminergic, and cholinergic systems. Transcriptomic enrichment analysis revealed associations with genes involved in mitochondrial function, synaptic signaling, and metabolic regulation. Together, these findings identify neural timescale disruption as a core pathophysiological feature of adolescent MDD, bridging macroscale dynamics with microcircuit and molecular architecture. INT may serve as a promising biomarker and mechanistic target for precision psychiatry in youth depression.

## Introduction

Adolescent major depressive disorder (MDD) has emerged as a significant public health concern due to its profound effects on the maturing brain and the unique clinical challenges it presents when compared to adult-onset depression. Adolescence represents a critical neurodevelopmental period characterized by rapid physiological, psychological, and social transitions, which contribute to pronounced interindividual variability in symptomatology and treatment responsiveness (Pines et al., 2023). A prominent characteristic of the adolescent brain is the ongoing cortical functional differentiation and the increasing complexity of neural processes. These neurodevelopmental changes play a critical role in the maturation of emotional regulation and cognitive functioning, thereby influencing responses to both pharmacological and psychotherapeutic interventions (Eiland & Romeo, 2013; Fournier et al., 2021). This intricate evolution of brain function may be associated with the temporal dynamics of neural activity, yet the dynamic neural representations of adolescent depression remain largely unknown.

The temporal characteristics of neural activity offer a window into this developmental functional differentiation, revealing a hierarchical organization of cortical timescales (Gao et al., 2020). Higher-order cortical areas tend to operate over longer timescales, enabling the integration of more complex emotional and cognitive information (Huntenburg et al., 2018; LeDoux & Brown, 2017; Singer, 2021). In contrast, primary sensory and motor cortices are typically governed by shorter timescales, facilitating rapid processing of external stimuli and motor responses (Hill et al., 2011; Zagha et al., 2013). Intermediate timescales, on the order of seconds, are likely involved in cognitive and affective operations such as working memory, decision-making, and emotional regulation (Lake et al., 2016; LeBlanc et al., 2015). These neural timescales, embedded within large-scale brain networks, are tightly linked to functional integrity; disruptions in regional timescales may impair global network coordination (Golesorkhi et al., 2021; Uhlhaas & Singer, 2012). Thus, analyzing neural dynamics through the lens of temporal hierarchy offers a promising framework for identifying spatially and temporally localized dysfunctions in psychiatric conditions.

In adolescent MDD, aberrations in cortical timescales may underlie core symptom domains. For instance, prolonged reaction times during emotional tasks may reflect deficits in affective processing or emotion regulation (Fournier et al., 2021; Heller & Casey, 2016; Park et al., 2024). Moreover, adolescent MDD is frequently accompanied by alterations in neuroplasticity, particularly maladaptive responses to chronic stress or adverse environmental inputs, which may exacerbate depressive symptoms and even contribute to suicidality (Dean & Keshavan, 2017; Ho & King, 2021). In healthy individuals, cortical timescales have been shown to be closely associated with multiple neurobiological substrates, including circuit-level parameters, cellular architecture, and genetic profiles (Dura-Bernal et al., 2024; Gjorgjieva et al., 2016; X.-H. Zhang et al., 2025). Therefore, elucidating cortical timescale abnormalities in adolescent depression may provide clinically relevant markers for symptom characterization and prognosis, while their microstructural underpinnings may inform biologically tractable therapeutic targets and guide etiological investigations.

Therefore, in the present study, we first identified abnormal patterns of cortical timescales by comparing cortical neuronal timescale differences between individuals with MDD and healthy controls (HC). Next, we employed a network lesioning approach to elucidate the impact of these aberrant regions on the global brain network architecture. Subsequently, we applied machine learning techniques to predict clinical symptoms of depression and examine their behavioral relevance. Finally, we further validated the circuit-level parameters, cellular characteristics, and genetic underpinnings associated with these aberrant cortical timescales.

## Results

### Demographic and clinical features

The demographic and clinical characteristics of the MDD and HC groups are summarized in S-table 1. Age was significantly different between groups (*t* = -5.80, *p* < 0.001), Age was higher in HC group, but there was no significant difference between groups in gender. For MDD patients, HAMD scores were 23.27 ± 3.64 (mean ± SD) and HAMA scores were 22.35 ± 6.42.

### The comparison between neural timescale of adolescent MDD and HC

Figure 1 illustrates the comparative analysis of neural intrinsic timescale (INT) derived from functional MRI (fMRI) blood oxygen level-dependent (BOLD) signals between adolescents with MDD and HC. Timescale calculation measures the temporal persistence of neural activity, characterizing how long neural activity maintains stability within specific brain regions. Longer INT indicates sustained neural activity and prolonged information integration, whereas shorter INT suggests rapid transitions in neural states (Figure 1a).

**Figure 1.**
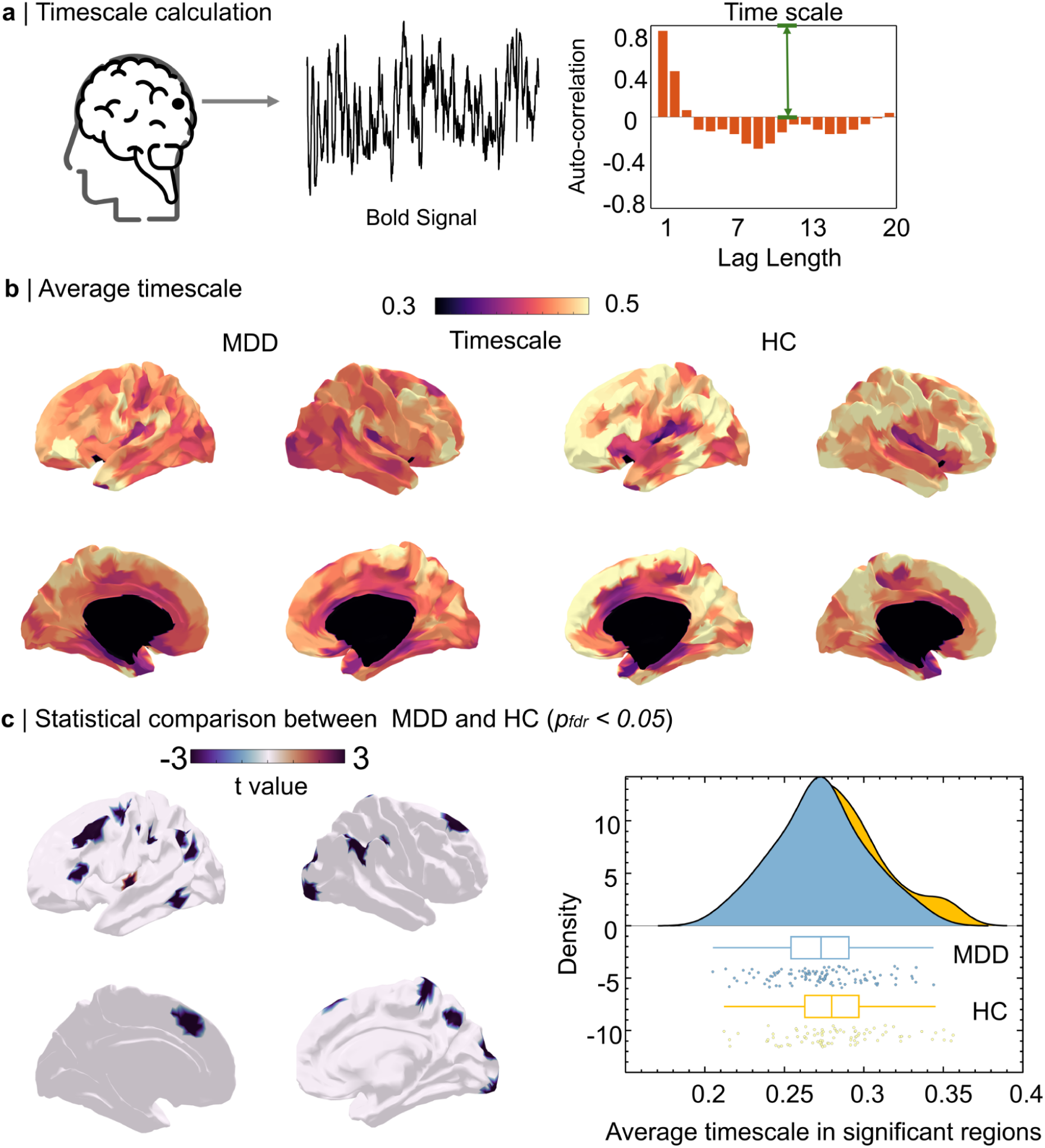
Comparison of neural timescale between MDD and HC. a. Timescale definition. Note that timescale was defined by the average of positive signal autocorrelation. b. Average timescale map in adolescent MDD and HC. c. Statistical comparison of timescale between adolescent MDD and HC (*p_fdr_* < 0.05) .

Spatial mapping of INT revealed that, in both MDD patients and HC, longer neural timescales predominantly occurred within frontal, parietal, and temporal cortices, indicating robust information integration in these areas. Conversely, sensory-motor, visual, and auditory regions exhibited shorter neural timescales, reflecting rapid neural processing (Figure 1b).

Critically, HC subjects consistently demonstrated elevated INT across most brain regions compared to adolescents diagnosed with MDD, who exhibited significantly reduced timescales. These findings indicate altered neural temporal dynamics associated with MDD (Figure 1b). Further statistical analysis confirmed significant group-level differences (p < 0.05, FDR-corrected), with MDD adolescents showing notably shorter INT in several key regions including the bilateral superior frontal gyri, left middle and inferior frontal gyri, left precentral and central sulcus, bilateral postcentral sulcus, left inferior temporal gyrus, bilateral inferior parietal lobules, right precuneus, and right superior/lateral occipital gyri. In contrast, the left temporoparietal junction (TPJ) exhibited a significantly longer INT in the MDD group compared to controls, suggesting differential regional disruptions of neural temporal dynamics associated with depression (Figure 1c).

### Network attack analysis for adolescent MDD and HC

Previous studies have demonstrated that timescale reflects the dynamic properties of the brain’s functional hierarchy. By removing brain regions with significantly altered timescales, researchers can elucidate the critical roles these areas play within the brain network, including their contributions to structural organization, information transfer efficiency, and overall network coherence.

Figure 2a illustrates the group-averaged functional connectomes for MDD and HC, computed from subject-specific Pearson correlation matrices at zero lag. As shown in Figure 2b, the removal of altered regions in the MDD group resulted in significantly higher modular disruption (two-sample *t*-test: *t* = –5.29, *p* < 0.001) and increased clustering coefficient disruption (*t* = –2.54, *p* < 0.001), compared to HCs. These metrics were computed by quantifying the difference in whole-brain modularity and clustering coefficient before and after node removal. Statistical comparisons controlled for age and sex as covariates. These findings suggest that in adolescent MDD, regions with aberrant intrinsic timescales play disproportionately central roles in maintaining the modular integrity and local segregation of the brain’s functional architecture. Their disruption may therefore contribute to the network-level disorganization associated with depressive psychopathology.

**Figure 2.**
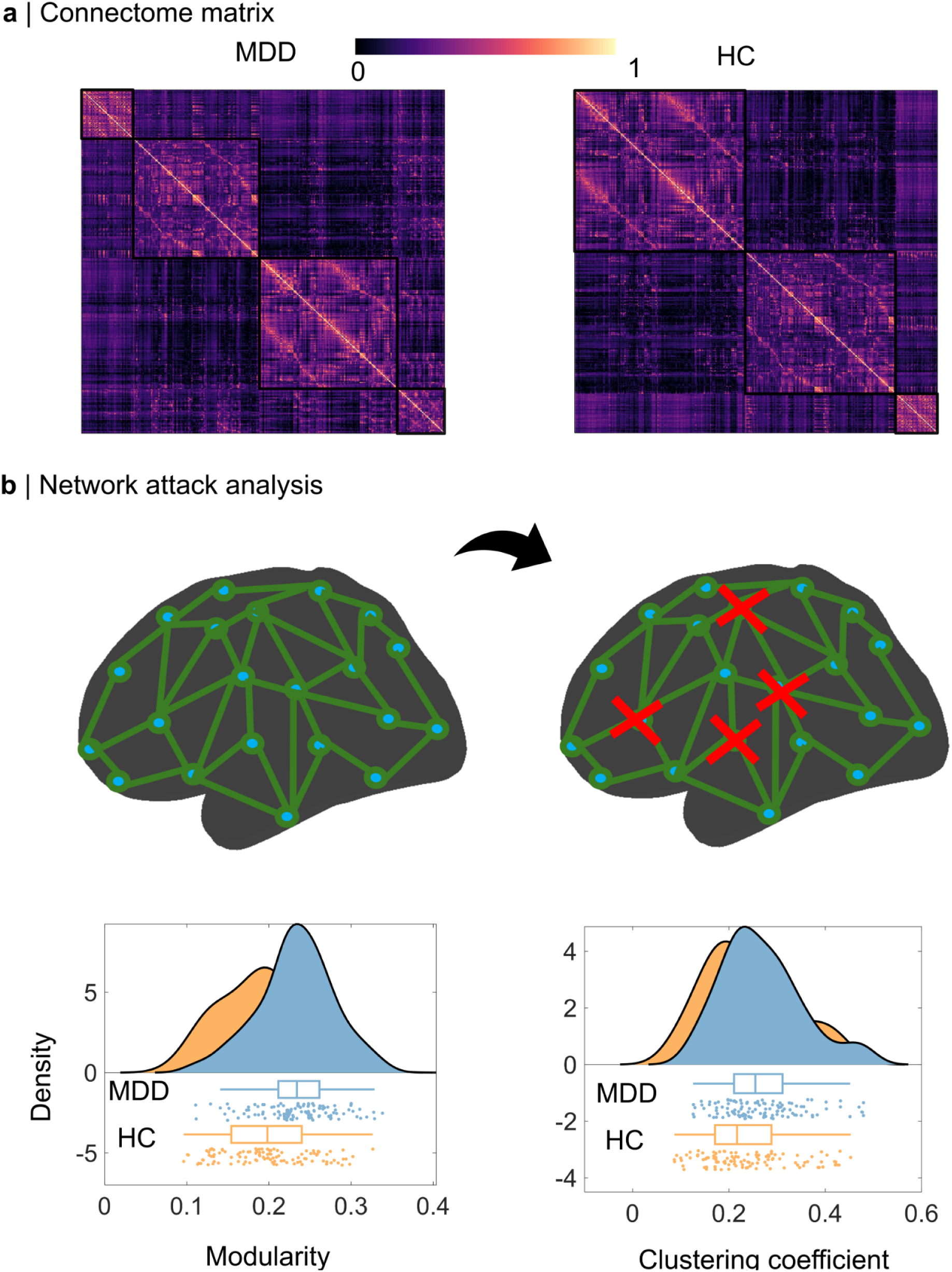
Network structure attack analysis in MDD and HC. a. The average functional connectome between MDD and HC. b. Networks attract analysis. We found adolescent MDD has significantly larger modular disruption (*t* = -5.29, *p* < 0.001) and clustering coefficient disruption (*t* = -2.54, *p* < 0.001) than HCs.

### Neural Timescale Predicts Clinical Symptom Severity in MDD

To evaluate the clinical relevance of altered neural timescales in MDD, we applied a multivariate regression framework to predict individual symptom severity from spatially distributed timescale patterns. Specifically, support vector regression (SVR) models were trained using neural timescale features that had shown significant group-level alterations between patients and healthy controls. A 10-fold cross-validation procedure, repeated 1,000 times to ensure robustness, was used to mitigate overfitting and to obtain stable estimates of predictive performance.

As shown in Figure 3a, neural timescales successfully predicted individual scores on both the Hamilton Anxiety Rating Scale (HAMA) and the Hamilton Depression Rating Scale (HAMD). Across subjects, predicted scores were significantly correlated with true clinical ratings (HAMA: *r =* 0.33, *p <* 0.05; HAMD: *r=* 0.32, *p <* 0.05), indicating that regional neural dynamics contain meaningful variance related to both anxiety and depression symptomatology.

**Figure 3.**
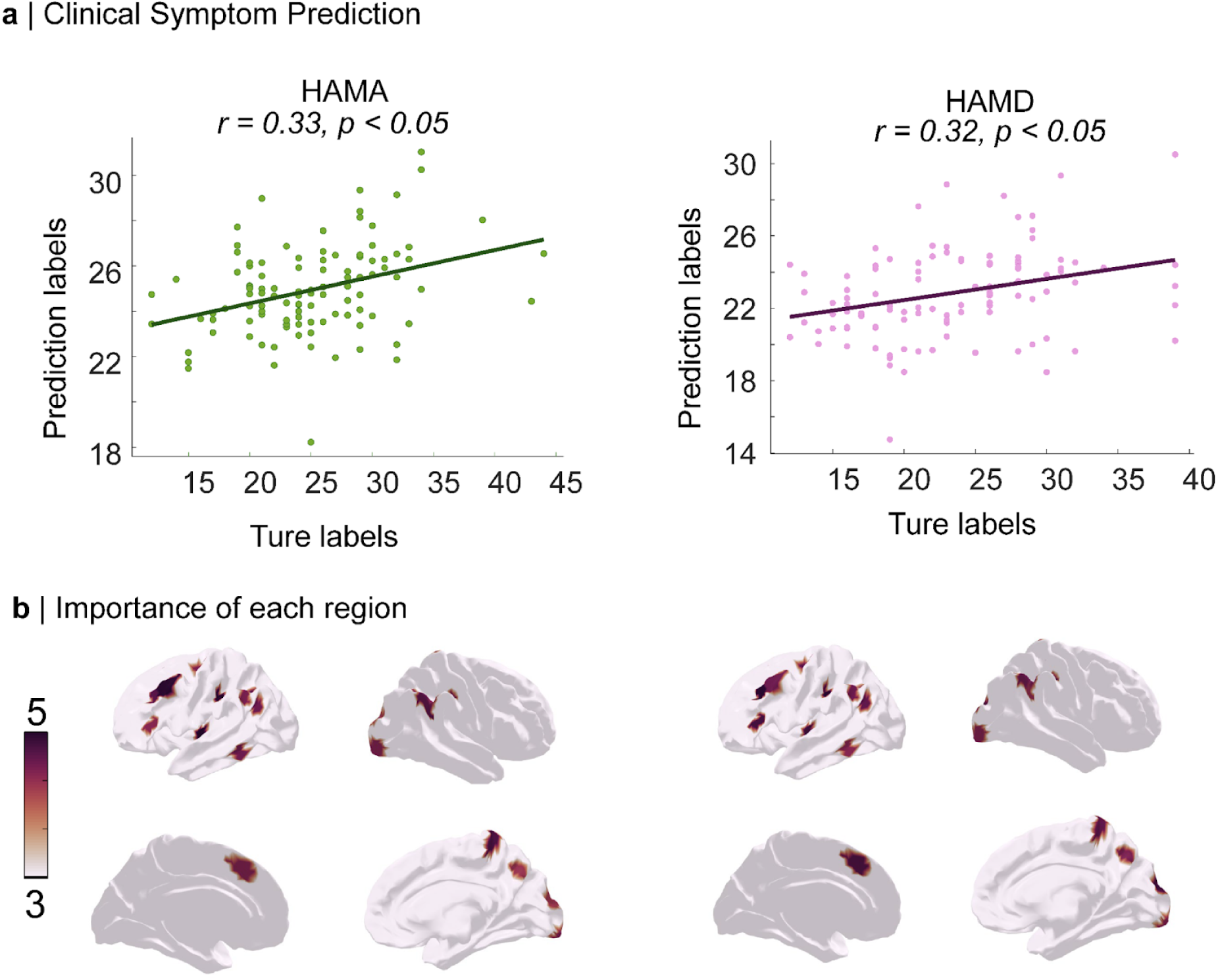
Timescale predicted clinical symptoms in MDD. a. Prediction of HAMA (*r* = 0.33, *p* < 0.05) and HAMD (*r* = 0.32, *p* < 0.05) via neural timescale. b. Importance of significant regions for HAMA and HAMD prediction.

To further dissect the spatial contributions underlying this predictive performance, we employed a random forest regression model to quantify feature importance across cortical regions (seen in Figure 3b). As depicted in Figure 3b, regions with the highest importance scores for symptom prediction included medial prefrontal cortex, posterior cingulate cortex, and anterior temporal lobe, overlapping with nodes of the default mode and salience networks. These regions are known to be implicated in affective regulation, self-referential processing, and emotional memory—core domains disrupted in MDD.

Collectively, these findings demonstrate that neural timescale alterations not only differentiate clinical populations but also serve as reliable predictors of symptom severity, providing biologically grounded markers for individualized assessment and potentially guiding targeted interventions.

### Clinical association with neural timescale

To investigate the clinical relevance of altered neural timescales, we first compared intrinsic timescale maps between individuals with suicide ideation (n = 30) and those without (n = 22). Voxel-wise analysis revealed significant group differences in specific cortical regions, with clusters showing reduced timescales in the suicide ideation group (*p_FDR_* < 0.05), suggesting that aberrant temporal dynamics may be associated with suicidality (Figure 4a).

**Figure 4.**
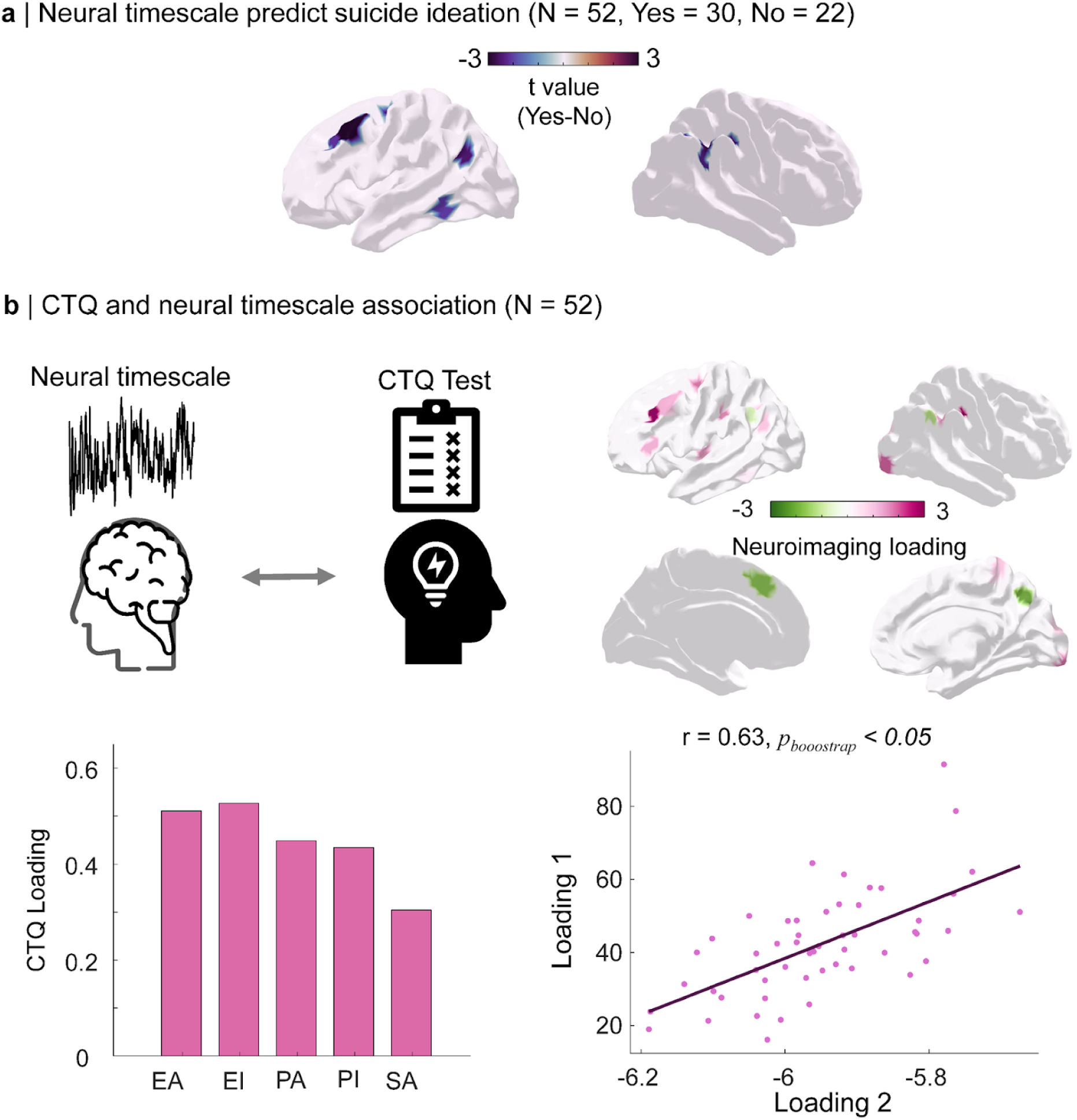
Clinical association with neural timescale. a. Comparison of timescale between suicide-paln patients and suicide-no-paln patients (*p_FDR_ <0.05*). b. Partial least squares analysis between various CTQ items and abnormal neural timescale (*r* = 0.63, *p_boostrap_ <0.05*).

Next, we examined whether these aberrant timescales were associated with early-life adversity, as assessed by the Childhood Trauma Questionnaire (CTQ). Using partial least squares (PLS) correlation analysis, we assessed the multivariate relationship between regional timescale alterations (X) and CTQ total and subscale scores (Y), including emotional abuse (CTQ-EA), emotional neglect (CTQ-EI), physical abuse (CTQ-PA), physical neglect (CTQ-PI), and sexual abuse (CTQ-SA). The first latent component derived from the PLS analysis exhibited a significant correlation between neural and behavioral loading patterns (r = 0.63, *p_bootstrap_* < 0.05; Figure 4b). Neuroimaging loadings were predominantly localized to prefrontal and temporoparietal regions, while behavioral loadings showed strongest contributions from emotional and physical neglect subscales.

These findings suggest that disruptions in neural temporal dynamics are not only associated with suicidal ideation but also significantly track with severity of early-life trauma, particularly neglect, thereby providing a potential neurobiological link between early adversity and current psychopathology.

### Regional Variation in Neural Timescale Is Associated with Circuit Parameters in a Biophysical Model

Using a biophysically informed parametric mean-field model (pMFM), we investigated how spatial variations in local cortical microcircuit properties relate to regional neural timescales. The pMFM integrates empirical structural connectivity, functional gradients, and cortical myeloarchitecture to simulate large-scale population dynamics, while allowing region-specific variation in intrinsic excitability and external drive.

Our modeling revealed that two circuit parameters—recurrent intra-regional excitation (denoted as www) and external input strength (III)—systematically vary with neural timescales across the cortex. Specifically, in HC, regional timescale estimates positively correlated with the strength of recurrent excitation (*r* = 0.23, *p_spin_* < 0.05; Figure 5, left panel), indicating that regions with longer intrinsic timescales tend to exhibit stronger local excitatory recurrence. In contrast, in individuals with major depressive disorder (MDD), timescales positively correlated with the magnitude of external input (*r* = 0.24, *p_spin_*< 0.05; Figure 5, right panel), suggesting a distinct influence of subcortical or global afferents in shaping temporal dynamics under pathological conditions.

**Figure 5.**
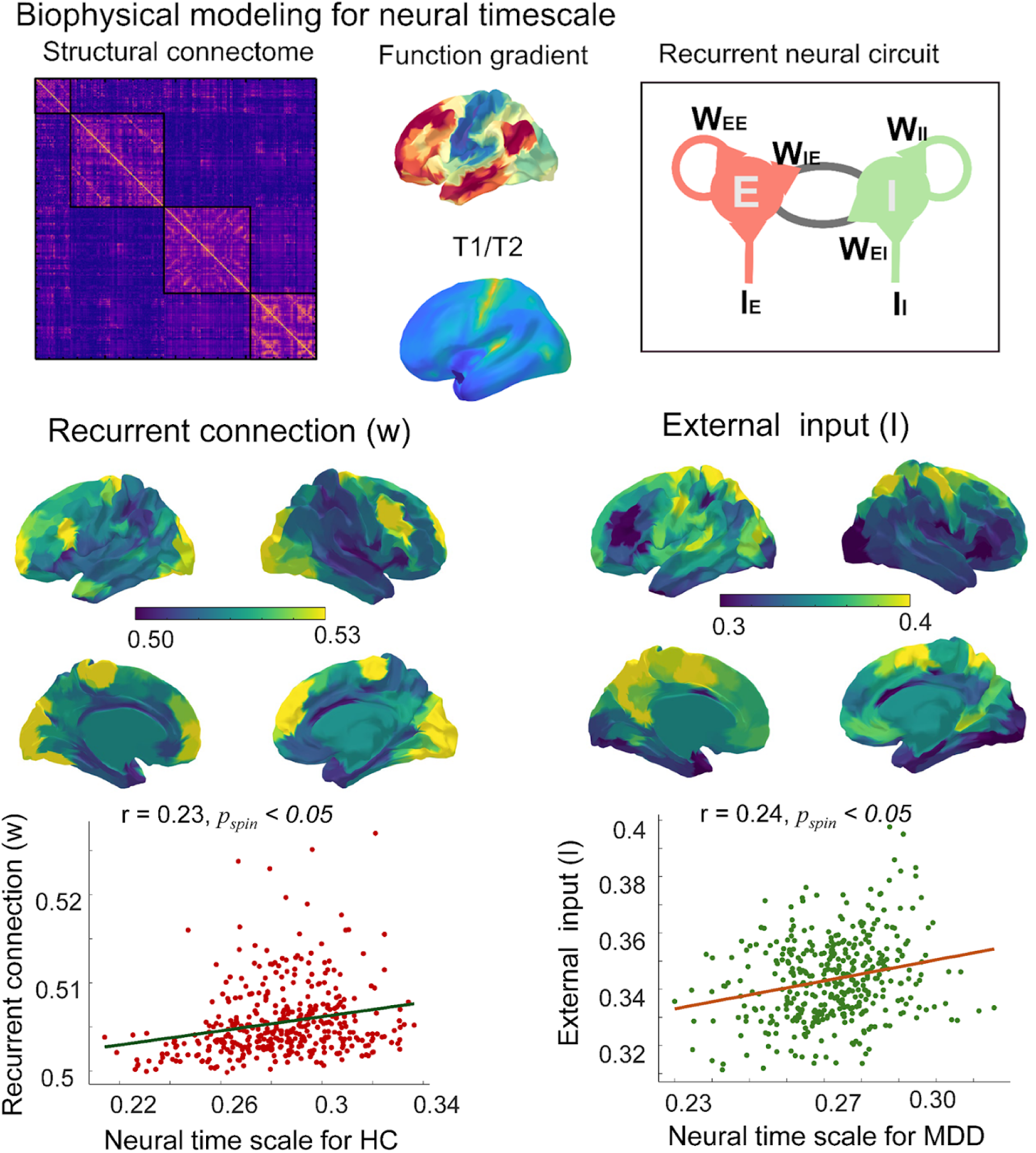
Biophysical modeling for neural timescale. Our results reveal the significant correlation between neural timescale with recurrent connection (*r* = 0.23, *p_spin_* < 0.05) and external input (*r* = 0.24, *p_spin_* < 0.05).

The spatial distribution of both www and III revealed a topographic gradient along the cortical mantle, closely aligning with known functional and anatomical hierarchies. Higher values of recurrent excitation and external input were localized to transmodal association cortices, including prefrontal and posterior parietal regions, consistent with their extended integration windows and functional roles.

Together, these findings support the hypothesis that regionally heterogeneous microcircuit mechanisms contribute to the large-scale organization of cortical temporal dynamics, and that this organization is differentially modulated in health and depression.

### Neurotransmitter Systems Are Topographically Coupled with Neural Timescale Alterations

To explore the neurochemical substrates of altered neural timescale organization in adolescent MDD, we examined the spatial correspondence between PET-derived neurotransmitter receptor/transporter (Hansen et al., 2022) and cortical timescale changes. Quantitative receptor density maps for major monoaminergic and cholinergic targets were aligned to cortical parcellations and correlated with regional timescale alterations. As illustrated in Figure 6, we observed significant positive spatial associations between abnormal neural timescale and the distribution of multiple neurotransmitter systems, including: 5-HT_1_A receptors (*r =* 0.31, *p_spin_ <* 0.05), Serotonin transporter (5-HTT) (*r =* 0.44, *p_spin_ <* 0.05), Dopamine transporter (DAT) (*r =* 0.36, *p_spin_ <* 0.05), Vesicular acetylcholine transporter (VAChT) (*r =* 0.42, *p_spin_ <* 0.05).

**Figure 6.**
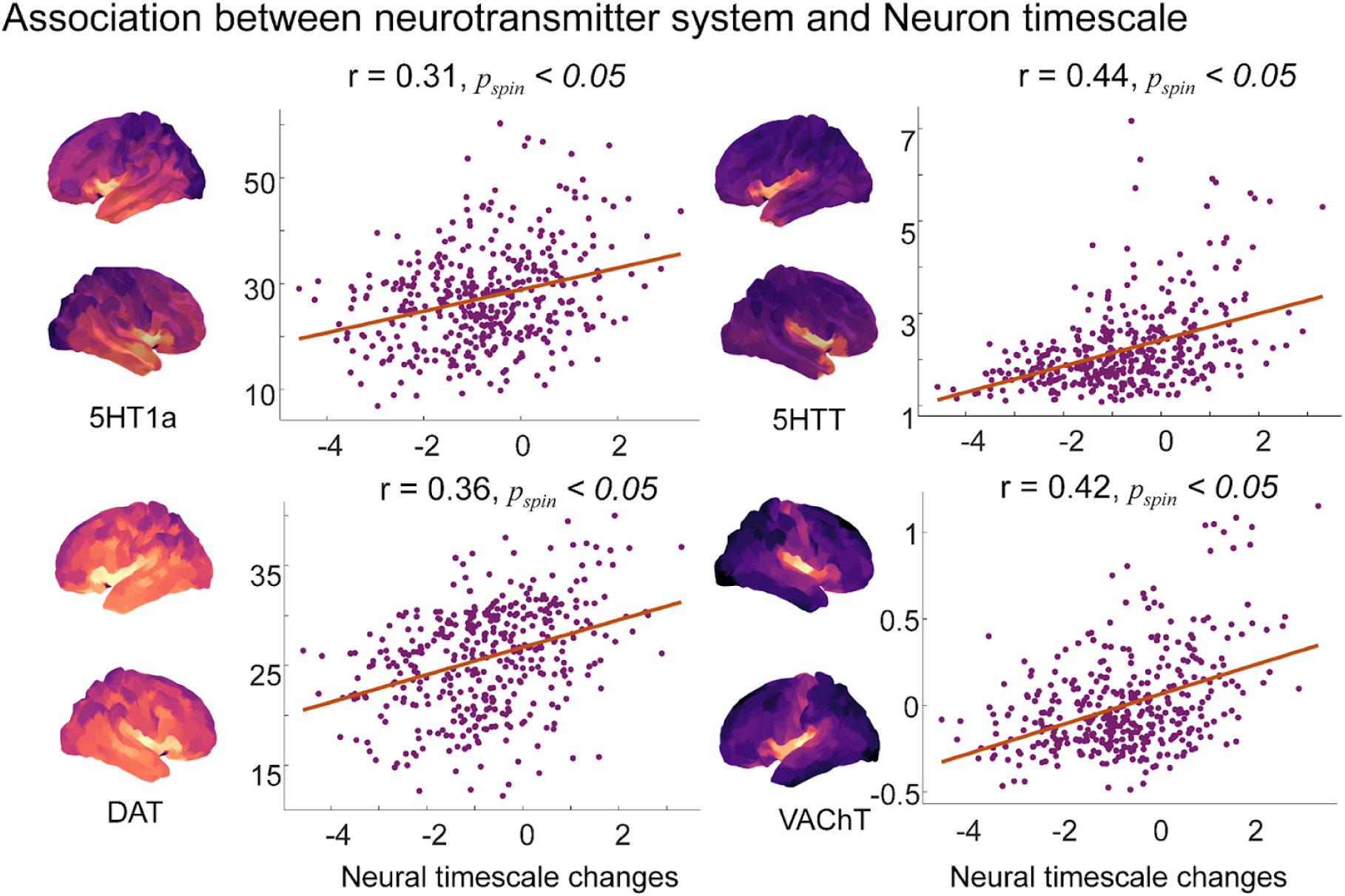
The correlation between neuron timescale with receptor maps, including 5HT1a (*r* = 0.31, *p_spin_* < 0.05), 5 HTT (*r* = 0.44, *p_spin_* < 0.05), DAT (*r* = 0.36, *p_spin_* < 0.05) and VAChT (*r* = 0.42, *p_spin_*< 0.05) .

These results were validated using a spatial permutation-based null model to account for cortical topographic autocorrelation. The observed correlations suggest that cortical regions with greater neurotransmitter receptor/transporter density also exhibit more pronounced timescale abnormalities, implicating neuromodulatory signaling—particularly serotonergic, dopaminergic, and cholinergic systems—in shaping temporal dynamics relevant to adolescent depression.

### Transcriptional Signatures and Biological Pathways Associated with Neural Timescale Abnormality

To uncover the molecular pathways underlying abnormal cortical timescales, we performed a spatially informed transcriptomic enrichment analysis. Gene expression data from the Allen Human Brain Atlas were matched to cortical parcellations using the abagen pipeline, and neural timescale statistical maps were correlated with regional mRNA expression patterns.

Gene sets exhibiting the strongest spatial correspondence to cortical timescale alterations were subjected to functional enrichment analysis using Metascape. As shown in Figure 7, significantly enriched biological processes and pathways (FDR-corrected) included: Aerobic respiration and respiratory electron transport, Mitochondrial organization, Inorganic ion transmembrane transport, Carboxylic acid and peptide metabolic processes, Synaptic signaling and neuronal system pathways, Regulation of protein localization and cellular component disassembly.

**Figure 7.**
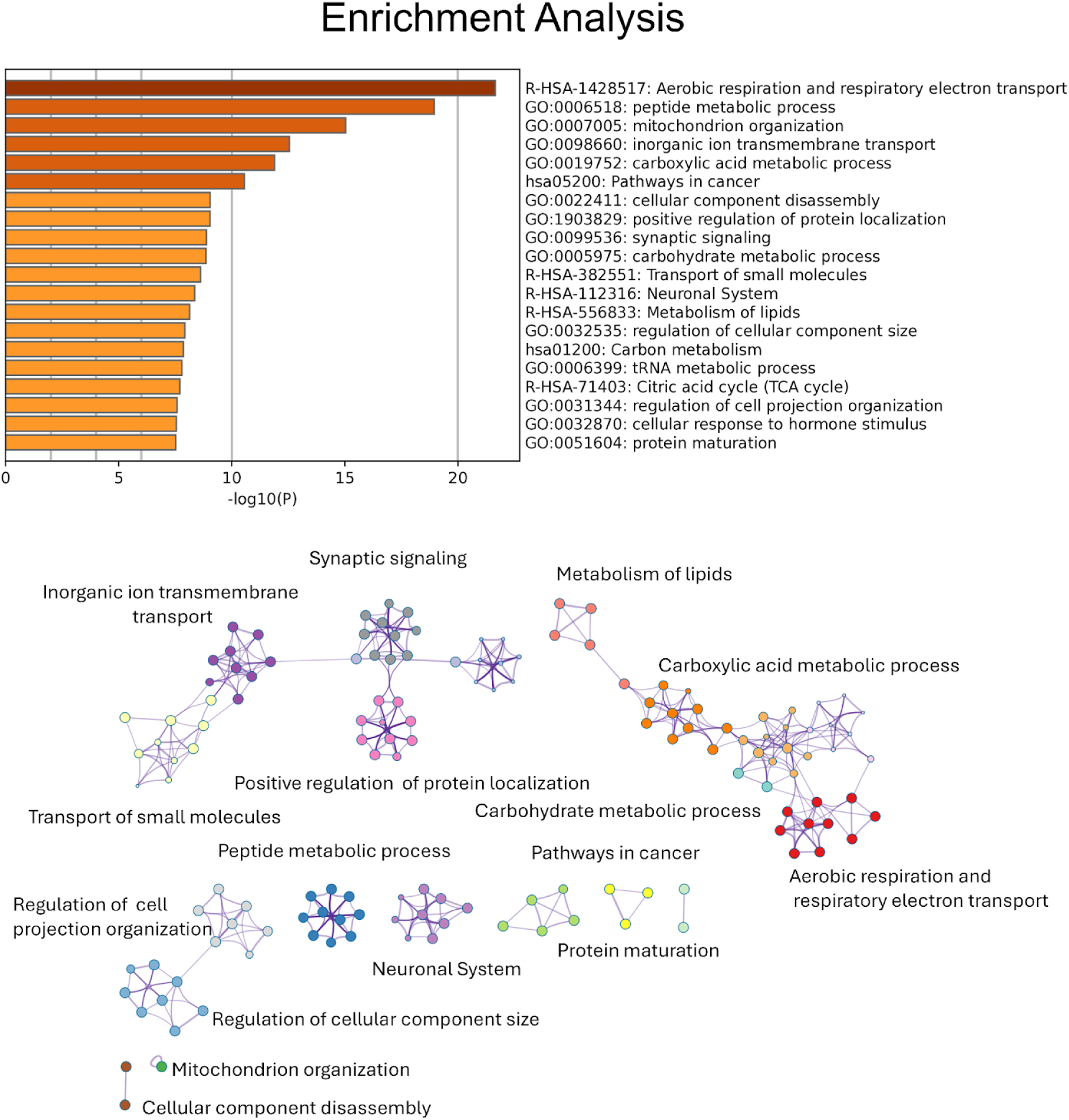
Enrichment analysis for abnormal neural timescale of adolescent MDD.

These findings indicate that energy metabolism, mitochondrial function, and synaptic signaling are central molecular themes associated with regional variations in neural timescale. Notably, the overlap with metabolic and synaptic pathways highlights the role of cellular bioenergetics and neurotransmission in sustaining temporal integration properties of cortical circuits—mechanisms potentially vulnerable in adolescent MDD.

## Discussion

This study found that adolescents with MDD exhibited widespread reductions in INT across multiple cortical regions, including the frontal, parietal, and sensorimotor areas, indicating impaired temporal stability of neural activity. In contrast, the left temporoparietal junction showed prolonged timescales, suggesting region-specific functional abnormalities. These alterations significantly disrupted the brain’s functional network architecture, reflected by reduced modularity and clustering. Neural timescales also predicted individual depression and anxiety severity, and were significantly associated with suicidal ideation and childhood trauma, particularly emotional and physical neglect. Mechanistically, INT was linked to regional variations in recurrent excitatory connectivity and external input strength, reflecting microcircuit-level dysfunction. Furthermore, altered timescales were closely associated with multiple neurotransmitter systems (5-HT_1_A, 5-HTT, DAT, VAChT) and enriched in molecular pathways related to mitochondrial metabolism and synaptic signaling, highlighting the role of neuromodulation and energy metabolism in the temporal dysregulation observed in adolescent MDD.

This study revealed that adolescents with MDD exhibited significantly reduced INT across widespread cortical regions, including the frontal, parietal, and sensorimotor cortices, suggesting impaired temporal integration and reduced stability of neural activity in these areas. Previous studies have shown that shorter timescales reflect faster yet less stable neural dynamics, indicating a diminished capacity for sustained information processing (Gao et al., 2020; Watanabe et al., 2019). Our findings extend this concept to adolescent depression, reinforcing the notion of disrupted temporal coordination in cortical networks. Interestingly, the left temporoparietal junction showed a significant prolongation of INT in the MDD group, in contrast to the generalized reduction observed elsewhere. The left temporoparietal junction is a critical node in the default mode and theory of mind networks, involved in self-referential thought, social cognition, and internal mentation (Buckner et al., 2008, 2009; Li et al., 2014; Vatansever et al., 2018). The increased timescale in this region may reflect hyper-stable neural dynamics, potentially contributing to maladaptive rumination and inward-directed attention (Hamilton et al., 2018; Whitfield-Gabrieli & Ford, 2012). This dissociation suggests an imbalance in the regulation of internal versus external attention in adolescent MDD (Diseth, 2005; Sommerfeldt et al., 2016). Future research may explore whether such regions represent potential targets for cognitive training or neuromodulatory interventions.

Moreover, network lesion analysis demonstrated that the removal of regions with abnormal timescales led to significant reductions in network modularity and clustering coefficients, underscoring their critical role in maintaining the brain’s functional architecture. These results align with prior evidence indicating impaired network segregation and integration in MDD (Fan et al., 2019; Kaiser et al., 2015; Luo et al., 2021). From a systems neuroscience perspective, these temporally altered regions may serve as functional hubs necessary for emotional regulation and executive control.

Machine learning analyses further demonstrated that neural timescale features reliably predicted individual depression (HAMD) and anxiety (HAMA) symptom scores, highlighting their clinical relevance. This suggests that INT metrics capture biologically meaningful variance in symptomatology and may serve as potential biomarkers for diagnosis or treatment monitoring. While prior research has linked dynamic functional features to affective symptoms (Marchitelli et al., 2022), our findings provide robust, cross-validated evidence in an adolescent population. Additionally, reduced timescales were significantly associated with suicidal ideation and early-life trauma, particularly emotional and physical neglect. This is consistent with previous work showing that adverse childhood experiences can lead to long-term alterations in brain structure and function (Herzog & Schmahl, 2018; Schaefer et al., 2022). Suicidal ideation has also been linked to dysfunctional default mode network activity (Malhi et al., 2020; S. Zhang et al., 2016), and our findings add a novel temporal dimension to this vulnerability, potentially enriching the neurobiological understanding of suicidality in depression.

At the circuit level, biophysically informed modeling showed that INT in healthy controls correlated positively with local recurrent excitatory connectivity, whereas in MDD patients, INT was more strongly associated with external input strength. This suggests a pathological shift toward increased reliance on bottom-up inputs and decreased intrinsic stability in MDD. These findings are consistent with the excitation-inhibition imbalance hypothesis of depression (Deco et al., 2024; Páscoa dos Santos & Verschure, 2022). Also, we further observed that regional INT alterations were closely associated with the spatial distribution of multiple neurotransmitter systems, including 5-HT_1_A receptors, 5-HTT, DAT, and VAChT. These neuromodulatory systems are central to mood regulation and are common targets of antidepressant treatments. The observed correlations suggest that disrupted neurotransmitter signaling may influence cortical temporal dynamics (Cipriani et al., 2018; Shine et al., 2019; Shinohara et al., 2019). Finally, transcriptomic enrichment analysis revealed that cortical regions with altered timescales were enriched for gene expression profiles related to mitochondrial function, aerobic respiration, synaptic signaling, and metabolic regulation. These pathways are critical for maintaining the energy demands and plasticity of cortical circuits, and have been implicated in the pathophysiology of depression (Manji et al., 2012; Picca et al., 2020; Scaini et al., 2016). Our findings suggest that the temporal dysregulation observed in adolescent MDD may have a molecular basis rooted in cellular bioenergetics and synaptic integrity. Future studies utilizing iPSC-derived neurons or brain organoids could validate these mechanisms and aid in the development of novel, metabolism-oriented therapeutic approaches.

## Limitation

Several limitations should be noted when interpreting the present findings. First, the study’s cross-sectional design limits causal inferences regarding the relationship between altered neural timescales and clinical symptoms or developmental trajectories. Longitudinal studies are needed to determine whether timescale alterations precede symptom onset or are a consequence of chronic psychopathology. Second, while our sample focused specifically on adolescents with MDD, heterogeneity in symptom profiles and medication histories may introduce variability not fully accounted for in our analyses. Third, although we integrated neuroimaging, biophysical modeling, and transcriptomic data, the resolution of PET receptor maps and gene expression atlases remains coarse and may not capture individual-level variability. Lastly, the generalizability of our findings to adult populations or other psychiatric conditions remains to be established.

## Conclusion

This study demonstrates that adolescents with MDD exhibit significant alterations in neural timescales, characterized by widespread reductions and region-specific prolongations, reflecting disrupted temporal integration and neural stability. These abnormalities are linked to impaired network organization, symptom severity, suicidal ideation, and childhood trauma, and are associated with circuit-level dysfunction, neuromodulatory systems, and mitochondrial-synaptic pathways. Our findings highlight neural timescale as a promising biomarker for adolescent depression with potential diagnostic and therapeutic implications.

## Method

### Ethics Statement

All study procedures adhered to the ethical guidelines of the Declaration of Helsinki and received approval from the Medical Ethics Committee of the First Affiliated Hospital of Chongqing Medical University. Prior to participation, written informed consent was obtained from the legal guardians of all participants. This study collected data from patients diagnosed with major depressive disorder (MDD) who sought treatment at the Department of Psychiatry and Psychology, Wuhan University People’s Hospital, between September 2021 and December 2023. Participant screening was conducted using the Chinese version of the Mini-International Neuropsychiatric Interview (MINI) to ensure diagnostic accuracy (Amorim, 2000).

The demographics of two datasets were shown in **Table 1**.

### Participants

This prospective study recruited adolescents diagnosed with depression and matched healthy controls (HCs) from August 2020 to July 2022. Participants were aged between 12 and 17 years. Adolescents with depression were enrolled from inpatient clinics at the Department of Psychiatry, First Affiliated Hospital of Chongqing Medical University, while HCs were recruited via community outreach.

Adolescent Depression Group: Participants experiencing their first depressive episode were included. Criteria included a Hamilton Depression Rating Scale (HAMD-24) score greater than 17, Han ethnicity, right-handedness, and no prior antidepressant treatment. Additionally, participants had not used psychotropic, anesthetic, sedative, hypnotic, or analgesic medications within one month before the study. Exclusion criteria included any history of manic or hypomanic episodes, neurological conditions (e.g., epilepsy, multiple sclerosis), significant physical illnesses (e.g., heart disease, cancer), other psychiatric diagnoses, borderline personality disorder, intracranial masses, family history of psychiatric disorders or self-harm, severe traumatic brain injury, substance or alcohol abuse or dependence, MRI contraindications, presence of metal implants or braces affecting imaging quality, and inability to cooperate with MRI procedures.

Healthy Controls (HCs): Healthy volunteers aged 12 to 17 years formed the control group and met identical exclusion criteria as the depression group. Additional criteria for HCs included a HAMD-24 score below 7, absence of major physical or psychiatric conditions, and no family history of psychiatric illness.

### Acquisition of Resting-State fMRI (rs-fMRI) Data

Resting-state functional MRI (rs-fMRI) data were acquired on a 3T GE Signa HDxt scanner (GE Healthcare, Chicago, IL) using an 8-channel head coil. Participants were instructed to remain awake with eyes closed, avoid cognitive tasks, and minimize head movements; comfort was optimized using foam pads and earplugs. None reported falling asleep during scanning. Echo-planar imaging (EPI) parameters included repetition time (TR)=2000 ms, echo time (TE)=40 ms, field of view (FOV)=240×240 mm, matrix=64×64, flip angle=90°, 33 slices, slice thickness/gap=4.0/0 mm, scan duration of 8 min, and a total of 240 volumes. Structural T1-weighted MRI images used for co-registration were acquired with TR=24 ms, TE=9 ms, FOV=240×240 mm, matrix=240×240, flip angle=90°, and slice thickness/gap=1.0/0 mm.

*Data preprocessing* Raw imaging data from all cohorts were initially acquired as DICOM files, subsequently reorganized into the Brain Imaging Data Structure (BIDS) format through HeuDiConv (version 0.13.1). Structural and functional neuroimaging data from both Bipolar groups underwent preprocessing utilizing fMRIPrep (Esteban et al., 2019), a pipeline operating on the Nipype framework (Gorgolewski et al., 2011). Structural preprocessing involved normalization of image intensity, extraction of brain tissue, segmentation into tissue classes, cortical surface reconstruction, and alignment into standard anatomical space. Functional images were corrected for head movement, adjusted for differences in slice acquisition timing, and spatially registered to structural reference images. Following preprocessing, the functional signals were partitioned using the Schaefer atlas 400. Confound regression was performed with Nilearn, adopting the simplified preprocessing pipeline described by (H.-T. Wang et al., 2024), incorporating high-pass temporal filtering, removal of motion artifacts (threshold was 2 mm), and physiological noise signals (e.g., from white matter and cerebrospinal fluid), linear detrending, and voxel-wise z-score standardization.

### Neural timescale definition

The Intrinsic neural timescales (INTs) reflect the temporal dynamics of information processing across brain regions, capturing regional differences in the speed at which neural information is integrated and processed. Here, we estimated INTs following the methodology outlined by Watanabe et al. (2019). For each participant, INTs were computed at the parcellation level using preprocessed resting-state fMRI data. The autocorrelation function (ACF) was first derived for each parcel, and the INT was quantified as the area under the ACF curve during its initial positive period. Group differences in INT were assessed using two-sample t-tests comparing adolescents diagnosed with MDD to healthy controls, with age and sex included as covariates in the model.

### Network attack analysis

Removing brain regions with significantly altered timescales in adolescents with MDD can provide deeper insights into the critical roles these regions play in network organization and global efficiency. In MDD, specific regions may exhibit aberrant timescales; by excluding these regions from network analyses, researchers can simulate the impact of their dysfunction on the overall brain network architecture. Therefore, we firstly generated subject-specific functional connectivity matrices based on pairwise Pearson correlations calculated at zero-lag from the processed regional time series. Furthermore, we quantified **modular disruption** and **clustering coefficient disruption** by comparing whole-brain modularity and clustering coefficient values before and after the removal of regions with significantly altered timescales. Finally, group differences in **modular disruption** and **clustering coefficient disruption** between adolescents with MDD and healthy controls were assessed using a *two-sample ttest*, with age and sex included as covariates.

### Recurrent Neural Circuit Modeling

To examine large-scale brain dynamics at the level of local microcircuits, we implemented a biophysically informed parametric mean-field model (pMFM), designed to simulate neural population activity across the whole brain while incorporating region-specific microcircuit heterogeneity. In contrast to prior models assuming uniform intrinsic properties across cortical areas, the pMFM permits key local neuronal parameters to systematically vary according to anatomical and functional cortical hierarchies, thus enabling biologically realistic simulations without excessive complexity.

Specifically, neural population dynamics within each cortical region were modeled as influenced by four primary factors: recurrent intra-regional excitation, inter-regional inputs modulated by empirically derived structural connectivity and scaled by a global coupling parameter, region-specific external inputs reflecting predominantly subcortical influences, and intrinsic neuronal noise. While global coupling was fixed uniformly across all regions, local parameters governing excitability, external drive, and noise were expressed as linear functions of two cortical gradients: the T1w/T2w myelin map and the principal gradient derived from resting-state functional connectivity in Brainspace toolbox (Vos de Wael et al., 2020).

### Clinical symptom prediction

To further elucidate the relationship between extracted neural timescale and clinical symptoms, as well as to validate the robust biological significance of specific episode patterns, we utilized a multivariate approach to predict clinical measures based on these patterns. More specifically, to predict individual symptom severity, including HAMD and HAMA scores, we trained support vector regression (SVR) models using as input the neural timescales previously identified as significantly altered between patients and healthy controls (Liu et al., 2021, 2024). Functional features were partitioned into training and test sets using a 10-fold cross-validation strategy to mitigate overfitting. In each fold, a linear-kernel SVR model (LIBSVM, https://www.csie.ntu.edu.tw/~cjlin/libsvm/, default parameters) was fitted on the training data and applied to the held-out test set. Predictive performance was quantified as the Pearson correlation between predicted and observed symptom scores. To ensure robustness, the entire cross-validation procedure was repeated 1,000 times, and averaged predictions across iterations were used in final analyses. In parallel, feature importance contributing to symptom prediction was assessed using a random forest regression model.

### Behavioral association with abnormal timescale

To examine the relationship between aberrant timescales and the Childhood Trauma Questionnaire (CTQ), we conducted a partial least squares (PLS) correlation analysis. Noted that CTQ comprises several subscales: Emotional Abuse (CTQ-EA), Emotional Neglect (CTQ-EI), Physical Abuse (CTQ-PA), Physical Neglect (CTQ-PI), and Sexual Abuse (CTQ-SA). The Total Score (CTQ-T) represents the overall severity of childhood trauma across these domains. The input variable set **X** consisted of regional brain areas showing abnormal intrinsic timescales, while the target variable set **Y** comprised the total CTQ scores and its subscales. To assess the statistical robustness of the primary PLS components, we employed a non-parametric permutation procedure. Specifically, brain features were held constant, while the rows of the target variables were randomly shuffled across participants. This procedure generated an empirical null distribution of Pearson correlations for the low-rank latent variables produced by the PLS analysis, representing the null hypothesis of random association between brain timescales and CTQ scores. For each iteration, the Pearson correlation between the permuted latent variables was recorded. The significance (p-value) of the observed component strength was then determined by comparing the original correlation to 1000 rho estimates derived from the null PLS models, with a significance threshold set at *p* < 0.05.

### Genetic analysis and enrichment analysis

To investigate the transcriptional underpinnings of large-scale network organization, we deconvolved cell type fractions from microarray expression data obtained from the Allen Human Brain Atlas (AHBA; http://human.brain-map.org/)(Shen et al., 2012). Gene expression data were derived from six neurotypical adult postmortem brains (mean age = 42.5 ± 13.4 years; 5 male, 1 female), comprising 3,702 spatially distinct cortical samples. To align transcriptomic and imaging data, we used the *abagen* toolbox (Markello et al., 2021) (https://github.com/netneurolab/abagen) to process microarray expression profiles and map them to Schaefer 400 atlas. We further correlated the neural timescale statistical maps with transcriptionally dysregulated genes in postmortem brain tissue of messenger RNA that are expressed most in cortical regions. To identify convergent or distinct biological pathways associated with functional reorganization, we performed gene enrichment analysis using Metascape (https://metascape.org) (Zhou et al., 2019), an integrative platform that synthesizes information from over 40 independent knowledge bases. Genes associated with the most prominent patterns of functional reorganization were submitted for analysis. Enrichment results were considered significant at a false discovery rate (FDR)–corrected threshold of *q* < 0.05, with significance further confirmed via comparison against a null model.

### Receptor maps

Neurotransmitter receptor and transporter densities were quantified using positron emission tomography (PET) tracer maps encompassing 18 molecular targets across nine major neurotransmitter systems, as curated by (Hansen et al., 2022) (https://github.com/netneurolab/hansen_receptors). These systems included dopamine (D_1_, D_2_, DAT), norepinephrine (NET), serotonin (5-HT_1_A, 5-HT_1_B, 5-HT_2_, 5-HT_4_, 5-HT_6_, 5-HTT), acetylcholine (α4β2, M_1_, VAChT), glutamate (mGluR_5_), GABA (GABA_a_), histamine (H_3_), cannabinoid (CB_1_), and opioid (MOR), consistent with prior neurochemical mapping efforts (Hansen et al., 2022) . PET images were nonlinearly registered to the MNI-ICBM 152 (2009c asymmetric) standard space and subsequently parcellated into the Schaefer-200 cortical atlas. For receptors or transporters with multiple PET maps derived from the same tracer (e.g., 5-HT_1_B, D_2_, VAChT), weighted averaging was applied to generate a single representative map per target, following the approach described by (Hansen et al., 2022).

### Null model

In the present study, we sought to quantify the topographic correspondence between neural timescale abnormality and other neurobiological features. To assess the statistical significance of these spatial associations while accounting for inherent spatial autocorrelation, we employed a spatial permutation-based null model (Markello & Misic, 2021). Specifically, receptor maps were first spatially aligned and correlated with neural timescale statistical map. To construct the null distribution, spherical rotations were applied to the cortical surface, systematically disrupting spatial alignment while preserving local spatial structure. Node values were reassigned based on the nearest rotated parcel, and this procedure was repeated 1,000 times. Rotations were initially applied to one hemisphere and then mirrored to the contralateral hemisphere to maintain interhemispheric symmetry. The empirical correlation was evaluated against the null distribution, with statistical significance defined by surpassing the *95th* percentile of the null correlations derived from both spatial and temporal permutation models.

## DATA AVAILABILITY

The clinical data could be accessed according to reasonable requests for corresponding authors. The raw fMRI data and MRI data for HCP was available on https://db.humanconnectome.org/. Heritability analyses were performed using Solar Eclipse 8.5.1b (https://www.solar-eclipse-genetics.org). Neuromap (https://netneurolab.github.io/neuromaps/usage.html), ENIGMA toolbox (https://enigma-toolbox.readthedocs.io/en/latest/pages.html).

## CODE AVAILABILITY

Code will be available on https://github.com/Laoma29/Publication_codes.

## ACKNOWLEDGMENTS

Xiaobo Liu is supported by the China Scholarship Council. Bin Wan is supported by International Max Planck Research School on Neuroscience of Communication: Function, Structure, and Plasticity (IMPRS NeuroCom), Graduate Academy Leipzig, and Mitacs Globalink Research Award. ZQL acknowledges support from the Fonds de Recherche du Qu\’ebec -- Nature et Technologies (FRQNT).This work was supported by in part by the Health of Hubei Province Scientific Research Project under Grant 2020Cfb512, and project of Mental Health Research Institute of Three Gorges University: YCXL-23-11.

## Supplementary Material

**Supplementary-Table 1.** Demography (HAMD means Hamilton Depression Rating Scale, HAMA means Hamilton Anxiety Scale, F means female, M means *male*)

| Group | Number | Age | Sex (F/M) | HAMD | HAMA |
| --- | --- | --- | --- | --- | --- |
| A-MDD | 110 | 14.18±1.51 | 35/75 | 23.27±3.64 | 22.35±6.42 |
| HC | 117 | 15.53±1.98 | 48/69 | ---- | ---- |

**Supplementary-Table 2.** Childhood Trauma Questionnaire (CTQ , CTQ-EA means Emotional Abuse, CTQ-EI means Emotional Neglect, CTQ-PA means Physical Abuse, CTQ-PI means Physical Neglect, CTQ-SA means Sexual Abuse, CTQ-T means Total Score), HAMA means Hamilton Anxiety Scale, F means female, M means *male*)

|  | CTQ-EA | CTQ-EI | CTQ-PA | CTQ-PI | CTQ-SA | CTQ-T |
| --- | --- | --- | --- | --- | --- | --- |
| Score | 12.75±4.88 | 15.88±5.56 | 7.5±3.50 | 10.40±4.07 | 6.08±2.50 | 52.65±14.92 |
| Number | 52 | 52 | 52 | 52 | 52 | 52 |

